# Navigating the Lipid Universe with LipidLibrarian: A Cross-Linked Database for Lipidomics Data Integration

**DOI:** 10.1101/2025.06.26.661298

**Authors:** Felix Niedermaier, Konstantinos Mechteridis, Konstantin Pelz, Vivian Würf, Nikolai Köhler, Josch K. Pauling

## Abstract

There are numerous public resources and guidelines available for lipidomics research, including standard nomenclatures, classification systems, and lipid databases. However, these resources are not always aligned with one another, making it difficult to find and compare information on the same lipid across different databases. To tackle these challenges we present LipidLibrarian, a lipid search engine that enables a combined search of all major lipid databases by aggregating the available information and presenting it in a unified manner. The three main sources of information that build the foundation of LipidLibrarian as a comprehensive search-engine are SwissLipids, LIPID MAPS and ALEX^123^. Furthermore, various secondary resources such as LION/web, LINEX, LipidLynxX, and Goslin were incorporated to enhance the results and conduct name and hierarchy conversions. LipidLibrarian is accessible via a user-friendly website, allowing the user to query lipids using their trivial names, shorthand notations, database identifiers, or their masses. Alternatively, LipidLibrarian can be accessed as a Python package for integration into high-throughput lipidomics pipelines. The output of a LipidLibrarian query is split into multiple categories, such as nomenclature, database identifiers, masses, adducts, fragments, ontology terms, and reactions. For each of these categories, LipidLibrarian aggregates the results from all databases and provides the source from which each value originates. This enables the user to quickly assess if the databases contain differing or conflicting information. In summary, LipidLibrarian provides an effortless, comprehensive and automated search for lipid information, thereby accelerating the research workflow and making it a meaningful tool for the scientific community.

## 1 Introduction

Over the years, the resolution of mass spectrometry instruments has improved, allowing us to identify lipids more accurately and with a better structural resolution [1]. While it was previously only possible to recognize lipid classes and sum species, modern instruments enable a more detailed determination of lipid structures, such as fatty acid chain lengths and number of double bonds. This allows for a classification of lipids into more precise subcategories. These subcategories are structured hierarchically, with the lower, more detailed levels containing the information from the broader, more general levels above. This means that each additional lower level not only adds new attributes to the lipids but also inherits the attributes from the higher levels. For instance, while the mass of the lipid is established at the sum species level, the positions of the double bonds are specified at the isomeric subspecies level, which then incorporates both the specific double bond positions and the previously determined mass from the higher level.

There are currently several lipid databases that specialize in different areas, such as physicochemical properties, fragmentation patterns, and reactions. A lipidomics research workflow often requires the acquisition of lipid information from multiple databases. However, the hierarchy systems used by the databases are not aligned to one another, making matching hierarchies a tedious task. In addition to the disparate hierarchies, the databases also use different notation styles to describe lipids. This makes it difficult to directly compare lipids with different notations and requires computational lipid parsers to check for equivalence. This issue could be solved by using cross-references between databases, but as they are not consistently provided for each entry and sometimes point to outdated records, they are currently insufficient for the development of robust computational workflows.

The divergence in hierarchy systems and notation styles between different lipid databases poses significant challenges to the collection and comparison of lipid data. Currently, these challenges in lipidomics research are addressed by manually aggregating and filtering the data from several databases for each individual lipid of interest. However, this is time-consuming, tedious and not suitable for high-throughput lipidomics workflows. A user-friendly solution overcoming these challenges is needed to accelerate and streamline lipidomics research. Taking on these challenges, we present LipidLibrarian, a lipid search engine that enables a combined search of all big lipid databases by aggregating the available information and presenting it in a unified manner. The three resources providing the necessary information to make LipidLibrarian into a well-rounded and extensive search-engine are SwissLipids [2], LIPID MAPS [3] and ALEX^123^ [4]. LIPID MAPS has a comprehensive database with a widely used hierarchy system, while SwissLipids provides Rhea [5] reactions for a lipid and ALEX^123^ contains comprehensive lipid fragment information. Additionally, we enriched the results with data from LION/web [6], and Rhea and Reactome [7] through LINEX [8]. To perform name and hierarchy conversions, we use LipidLynxX [9] and Goslin [10]. LipidLibrarian is developed as an open-source application with a focus on extensibility to easily integrate new resources.

LipidLibrarian can be accessed through a user-friendly website, allowing users to search for lipids by their trivial names, shorthand notations, database identifiers or masses. Furthermore, the tool can be integrated into high-throughput lipidomics pipelines as a Python package. The software is intended for a wide range of users, from beginners to experts in computational biology or lipidomics research. The output of LipidLibrarian is presented in an intuitive results page, consisting of multiple categories that hold information on the queried lipid: nomenclature, database identifiers, masses, adducts, fragments, lipid ontology terms, and reactions. For each of these categories, LipidLibrarian aggregates the results from all databases into a consistent layout, such that it is traceable from which source each value originates. This layout also enables the user to quickly assess if the databases contain differing or conflicting information.

In this paper, we will present the methods used by LipidLibrarian to efficiently search, acquire, align, and merge lipid information across multiple databases. We will introduce and discuss the two versions of LipidLibrarian, the Python package and the corresponding web interface. Possible applications of both versions will be demonstrated using different lipid examples to provide the reader with a comprehensive understanding of the tool’s capabilities.

## 2 Materials & Methods

### 2.1 Information Sources

In order for LipidLibrarian to acquire comprehensive information on a given lipid, it makes use of several primary and secondary information sources. The primary information sources, consist of the the three lipid databases ALEX^123^, LIPID MAPS and SwissLipids. The secondary information sources, LION/web (Lipid Ontology) and LINEX are used to enrich the results acquired from the primary sources

#### 2.1.1 ALEX^123^

Pauling et al. [4] propose a “common nomenclature for annotation of lipid fragment ions” which can be used to query a web tool called ALEX^123^ lipid calculator, available at http://alex123.info. This tool covers approximately 430,000 lipid molecules from 47 lipid classes mainly from the lipid categories: fatty acyls, glycerolipids, glycerophospholipids, prenol lipids, sphingolipids, and sterol lipids.

Although ALEX^123^ contains data from MS1, MS2 and MS3 experiments, LipidLibrarian only uses MS2 queries to extract lipid fragmentation information. For performance reasons the MS2 data was extracted from the web tool using a script that scrapes all available lipids, and by querying the website using all possible *m/z* (mass-to-charge) ratio and tolerance values. The results are combined into a database containing approximately 295,000 molecular species and 8,600 sum species.

#### 2.1.2 LIPID MAPS

The second primary resource is LIPID MAPS (LIPID Metabolites And Pathways Strategy), which is a consortium founded in 2003 to provide resources for lipids and aid the international lipid research community. The consortium divides lipids into the following eight categories, a classification that is accepted by most researchers today: fatty acyls, glycerolipids, glycerophospholipids, sphingolipids, sterol lipids, prenol lipids, saccharolipids, and polyketides. [11]

The Lipid MAPS Structure Database (LMSD) contains, as of June 2025, 49,761 lipid entries, originating from the LIPID MAPS Consortium’s core laboratories and partners. These lipids are identified experimentally, computationally using lipid classes or are manually curated from LIPID BANK, LIPIDDAT and other sources based on biological relevance [3].

Each lipid has a unique LMID which is 12 or 14 characters long and follows a specific nomenclature and hierarchy defined by LIPID MAPS [11, 12, 13].

In order to acquire lipid data from LIPID MAPS we use the provided REST API, further described on the consortium’s website [14], which returns us chemical data, nomenclature and hierarchy information, and structure identifiers.

#### 2.1.3 SwissLipids

SwissLipids [2] is a knowledge-database, developed by the SIB Swiss Institute of Bioinformatics. As of June 2025 it consists of 779,249 lipid species and 7170 distinct pieces of curated evidence from 1580 peer-reviewed publications. It was created by experts curating lipids from different resources like ChEBI [15], Rhea [5], or UniProt [16] and enriched by a library of computed theoretical lipid structures.

All lipids contain standard nomenclature, cheminformatics descriptors, ontological information like UniProt [16], ChEBI [15], or GeneOntology [17, 18], formula, and mass. This information is often supplemented with cross-links to other databases. Every lipid also contains a unique identifier which is 13 characters long and starts with the term “SLM:”. It uses the nomenclature and the classification scheme of LIPID MAPS [19], but has a separate hierarchy system [2].

To query SwissLipids we used the Rest API provided by the database, which can be found on the SwissLipids website [20] and allows to extract chemical data, structure identifiers, reactions and cross references to other databases.

#### 2.1.4 Secondary Information Sources

Unlike primary information sources, secondary sources only contain data for a certain set of lipids and often accept specific inputs, such as the lipid class abbreviation. These sources are only queried after all the baseline data has been collected from the primary sources. Following three secondary sources are used by LipidLibrarian to enrich the retrieved lipids:

First, the **LINEX2** Python package [8], which combines all the reactions from both RHEA [5] and Reactome [7]. By querying this source, we can provide information about which other lipid classes our returned lipids react with. This gives users a broader understanding of their query results. Second, the lipid ontology database, developed as a part of the **LION/web** application. It provides a substantial dataset of lipids mapped to properties such as chemical characteristics, lateral diffusion, and transition temperatures as ontology terms [6]. Finally, we implemented the **LipidLibrarianAPI**, which is used for minor offline processing and data completion purposes.

### 2.2 Lipid Nomenclature

In order to describe or compare lipids, a suitable grammar must be defined. The demands on this nomenclature are completeness, consistency, human and at the same time machine interpretability, and above all no ambiguity where none is wanted.

Lipids can, of course be referred to, like all molecules, by their sum formula, IUPAC name [21, 22], SMILES [23] or InChI [24]. All of these nomenclatures are not ideal for describing lipids when presenting them in a database or discussing them in a publication. They are either not very human readable and interpretable, or they generate quite long and repetitive patterns for simple common features of lipids. A phosphocoline head group, for example, is described by 25 characters in the IUPAC nomenclature and 55 characters as a SMILES string. A simple n-carbon fatty acid which is at least *n* characters as a SMILES string or 2*n* as an InChI string. Furthermore, we may not always know the exact structure for all features like double bond location and order of fatty acids, and still need to be able to compare and discuss these less strongly defined molecule classes.

The Shorthand Notation for Lipids [11, 25, 26] is designed to accommodate for special Lipidomics needs. It is able to express broad definitions of lipid species and classes, while also allowing finegrained control over details and changes within the definition, for example the order of fatty acids or location of double bonds for isomeric subspecies. At the same time the notation keeps the focus on the essential features, and therefore offers the necessary brevity.

The shorthand notation for a phospholipid sum species consists of an abbreviation for the phospholipid class, followed by number of C-atoms and the number of double bond equivalents, i.e., PS 36:4 [26]. If there is more than one fatty acid connected to the headgroup and the exact carbon chain length and double bond number of each of them is known, the shorthand notation expresses them separated by slashes, i.e. TG 12:0/18:2/22:1. If the sn-positions are unknown, the fatty acids are separated by underscores instead of slashes.

The core idea and semantic of the Shorthand Notation is generally accepted within the community, however multiple standards and syntaxes between research groups and databases have evolved. The matter was further complicated by the lack of a standardized nomenclature for lipid fragments until the Pauling, J. K. et al. paper was published in 2017 [4]. Due to these sources of ambiguity, details are different and it is not trivial to match one naming variant to another.

#### 2.2.1 Nomenclature Parsing

LipidLibrarian’s ability to parse lipid names and interpret nomenclature is one of its core features. LipidLibrarian is able to mediate between databases with different naming schemes and combine information found on the same lipid, even if found under different names. There are already tools available to assist in the parsing of lipid handling nomenclature, including REST APIs, locally executable applications and libraries. This section discusses a selection of relevant ones that were considered throughout the LipidLibrarian development process.

LipidLynxX implements a complex regex parsing system and is able to convert nomenclature from the databases HMDB, LIPID MAPS LMSD and COMP DB, LipidHome, RefMet, SwissLipids, as well as various shorthand notations (e.g., space-separated like PC 16:0_18:2 and bracketed like PC(16:0_18:2)), trivial names and common abbreviations in literature to an internal representation, which can then be output as shorthand notation using either the space or bracket format [9].

The newer lipid nomenclature parsing tool Goslin unifies nomenclature between databases and provides implementations to convert all of the dialects into a standardized name according to the lipid shorthand nomenclature, to which we previously referred to as the LIPID MAPS flavour [27, 10]. The Goslin grammar is defined using ANTLR 4, a language recognition syntax independent of any specific programming language. There are implementations of Goslin in various programming languages, e.g. pygoslin written in Python [28] or a web interface with API access [29].

LipidLynxX and Goslin interchangeably help LipidLibrarian to convert lipid notation between the shorthand notation dialects, and conversions between detail levels, e.g. determining the sum species of an isomeric subspecies.

The liputils Python package is a lightweight lipidomics analysis tool primarily used for handling and quantifying lipids in Python after a mass spectrometry analysis [30]. LipidLibrarian integrates liputils because it contains a number of handy functions for extracting lipid properties from their shorthand representations.

### 2.3 LipidLibrarian Python Library

At the heart of LipidLibrarian is the Python package. It bundles all the code for requesting information about lipids from the APIs and provides the experienced user with the ability to automatically request all of these APIs and combine and complement their results in a logical way. This code library is not used directly by the user, but requires an executable in which it is embedded. In this project we provide two implementations. One is a command-line program that outputs the lipids found as text or files, and the other is a web application that is accessed via the user’s browser and displays the results. Both implementations are described in more detail in the results chapter.

#### 2.3.1 Workflow

The typical workflow when using LipidLibrarian can be subdivided into five steps as represented in Figure 1:

**Figure 1.**
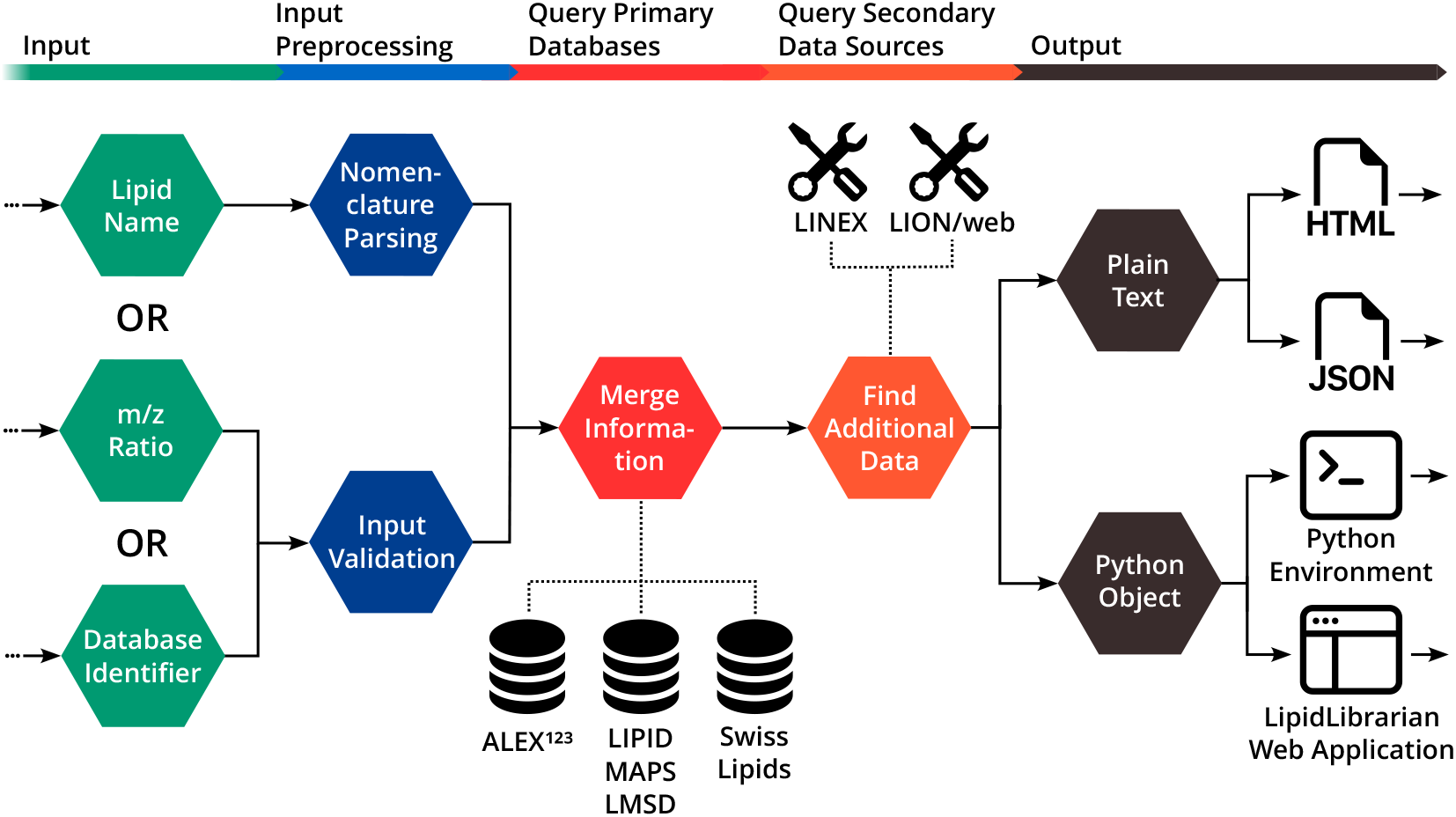
Conceptual workflow of LipidLibrarian. The user starts the workflow by either supplying a lipid name, an *m/z* (mass-to-charge) ratio, or a database identifier. LipidLibrarian processes the input, dynamically aggregates information from online and offline databases and provides the output data in the desired format.

1. **User Input Choices**: In the first step, users can select one of three input types to use for their query. One possible input for a search is the lipid name, which is a string containing the shorthand name of a lipid. Inspired by the hierarchy system of LIPID MAPS and SwissLipids, the user is able to submit a lipid name in one of four hierarchy levels, which are: Sum Species, Molecular Subspecies, Structural Subspecies and Isomeric Subspecies (example in Table 1). Another input option is to specify a *m/z* (mass-to-charge) ratio and a tolerance range in which the extracted lipids should be included in. Lastly, LipidLibrarian also supports database identifiers from LIPID MAPS, SwissLipids, and ChEBI as an input type.
2. **Preprocessing**: Depending on the input choice of the user the preprocessing step is divided into two parts. The first part, nomenclature parsing, is only used if a lipid name was provided as input. Here the input string is parsed to determine the lipid class and the hierarchy level of the lipid and to check whether the lipid name is valid. In the second part where an *m/z* ratio or database identifier was provided, the values are only checked if they are of the appropriate data type and have the correct formatting.
3. **Query Primary Databases**: After the preprocessing is complete the input values are queried in the three databases ALEX^123^, LIPID MAPS LMSD and SwissLipids. The acquired information of all three databases is merged within a lipid Python object. If there is a disagreement about a feature between two data sources, LipidLibrarian will retain both versions, including information about what the query was and which database produced the value, so that the user can decide how to resolve the conflict. To obtain even further information about our input, LipidLibrarian offers a setting to run this step and query all databases again with the additional information gained on the just completed run.
4. **Query Secondary Databases**: Similar to the previous step the secondary databases LINEX, LION/web (Lipid Ontology) and the internal functions in LipidLibrarianAPI are queried, to acquire additional information on our lipids.
5. **Output**: Finally, our lipids are returned as Python objects the user can use to perform further downstream analysis. The lipid information can also be returned as text files in HTML or JSON format.

**Table 1:** The hierarchy levels of LipidLibrarian are illustrated using a phosphatidylcholine as an example.

| Level | Value |
| --- | --- |
| Sum Species | PE O-42:7 |
| Molecular Subspecies | PE O-18:1_24:6 |
| Structural Subspecies | PE O-18:1/24:6 |
| Isomeric Subspecies | PE O-18:1(13Z)/24:6(6Z,9Z,12Z,15Z,18Z,21Z) |

### 2.4 LipidLibrarian Command Line Interface Technology

The command-line interface is a thin Python wrapper around the main library. It offers the same options as when importing the library in a Python environment and outputs the results to the terminal or as text files.

### 2.5 LipidLibrarianWeb Application Technology

The LipidLibrarian web application is implemented with a modern stack of web development technologies consisting of the Angular and Django web development frameworks and a PostgreSQL database. The three components, frontend, backend, and database, are divided into separate containers that communicate with each other through a virtual network (see Figure 2). The Django backend executes the LipidLibrarian code, which retrieves lipids and stores them in the Postgres database. To request a lipid, users interact with the Angular frontend which in turn retrieves information from the backend using an Application Programming Interface using Representational State Transfer (REST API). Finally, the lipids are displayed on the website.

**Figure 2.**
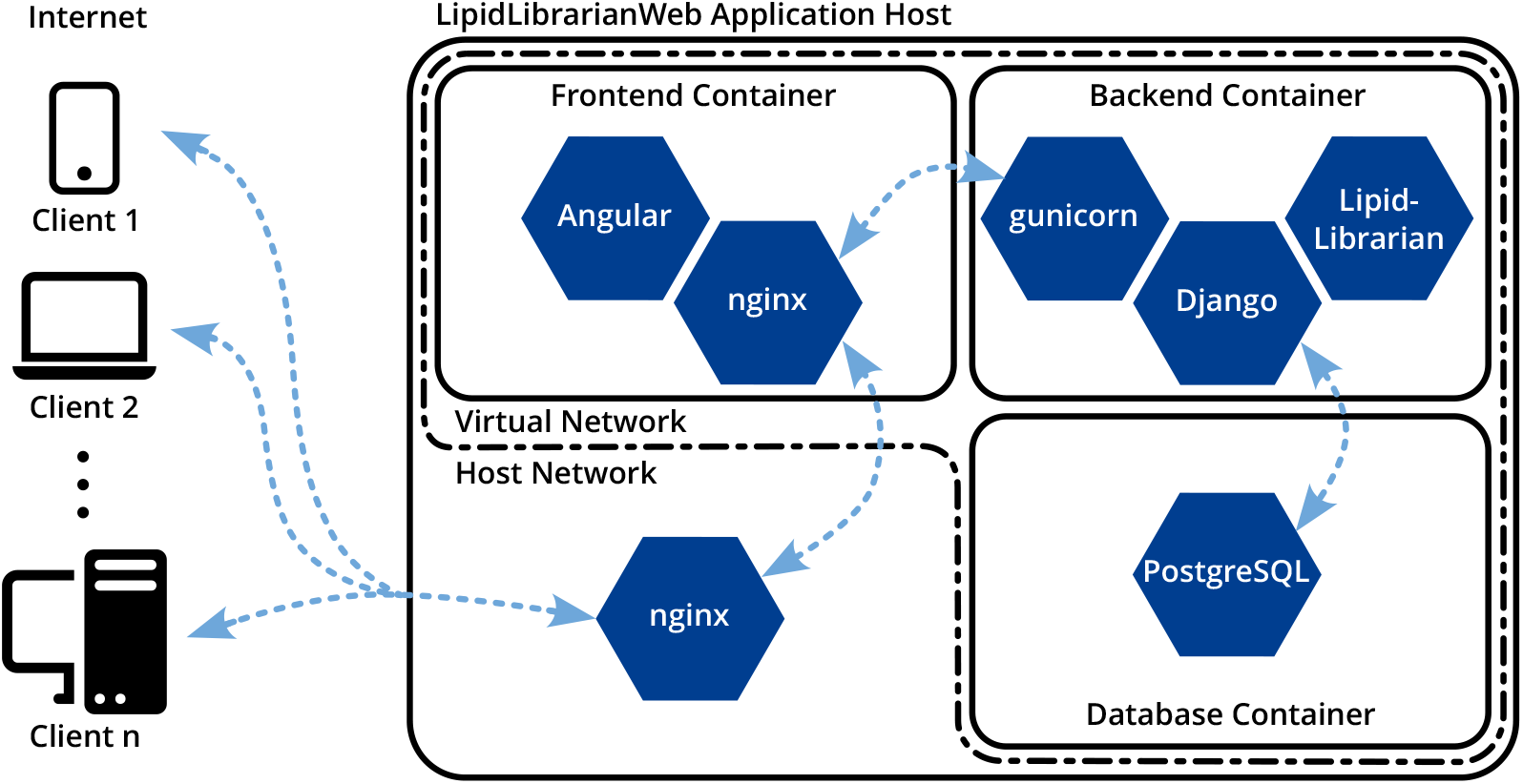
Overview of the LipidLibrarianWeb architecture. LipidLibrarianWeb is accessed from the client devices via an HTTPs connection in their browser. The requests are handled by a web server (nginx) on the application host, which forwards them to the single point of entry: the web server (nginx) of the Frontend Container that serves the website and proxies the REST API requests to minimize the attack surface. The backend container wraps the LipidLibrarian python package and handles the connection to the database for caching queries and results.

## 3 Results

LipidLibrarian is currently available as two implementations, as a command-line interface and as a web application that displays the obtained information on a user-friendly website. They both use the LipidLibrarian Python library to access the shared code. In addition, it is easy to import the LipidLibrarian library into a Python environment, such as a Jupyter notebook, and use it in that environment.

### 3.1 LipidLibrarian Command Line Interface

The command line interface is intended for programmers, bioinformaticians and for testing the library. It is suitable for use in pipelines, as it has support for piped input and output in UNIX shell environments, and can handle large batch requests from CSV files. When running LipidLibrarian from the command line, the user has several options, including whether to save the information from the lipids acquired as an HTML or JSON file, which databases to inand exclude from the query, and how often to repeat the query process with the new found information.

### 3.2 LipidLibrarianWeb

Since bioinformaticians, computational biologists and chemists who are able to understand and write Python code or know their way around the command line are not the only target group for LipidLibrarian, LipidLibrarianWeb, an easy to use web application for querying LipidLibrarian was developed. With LipidLibrarianWeb any interested user may search for lipid trivial names and more experienced users may query the application with *m/z* ratios and tolerances from their experiments conveniently from the web browser.

#### 3.2.1 Workflow

The LipidLibrarianWeb workflow begins when a user initiates a query through the website. The detailed workflow is depicted in Figure 3 and starts with the user filling in one of the three query options, lipid name, *m/z* ratio, or database identifier, and then submitting the request. There is a search field always accessible in the navigation bar at the top of the website so the user can access the field from all sub-pages of the site and perform a quick search. Alongside the basic search field, there is an advanced search form on the main page. Here the user can enter either a search term or a *m/z* ratio with a tolerance range in which the extracted lipids should be contained. In the case of a *m/z* search, the user can also select which positive and negative adducts are to be used for the search. Additionally, the advanced search allows the user to select the databases in which LipidLibrarian should search for the particular input. After filling out the required input fields, the user submits the advanced query to start the search.

**Figure 3.**
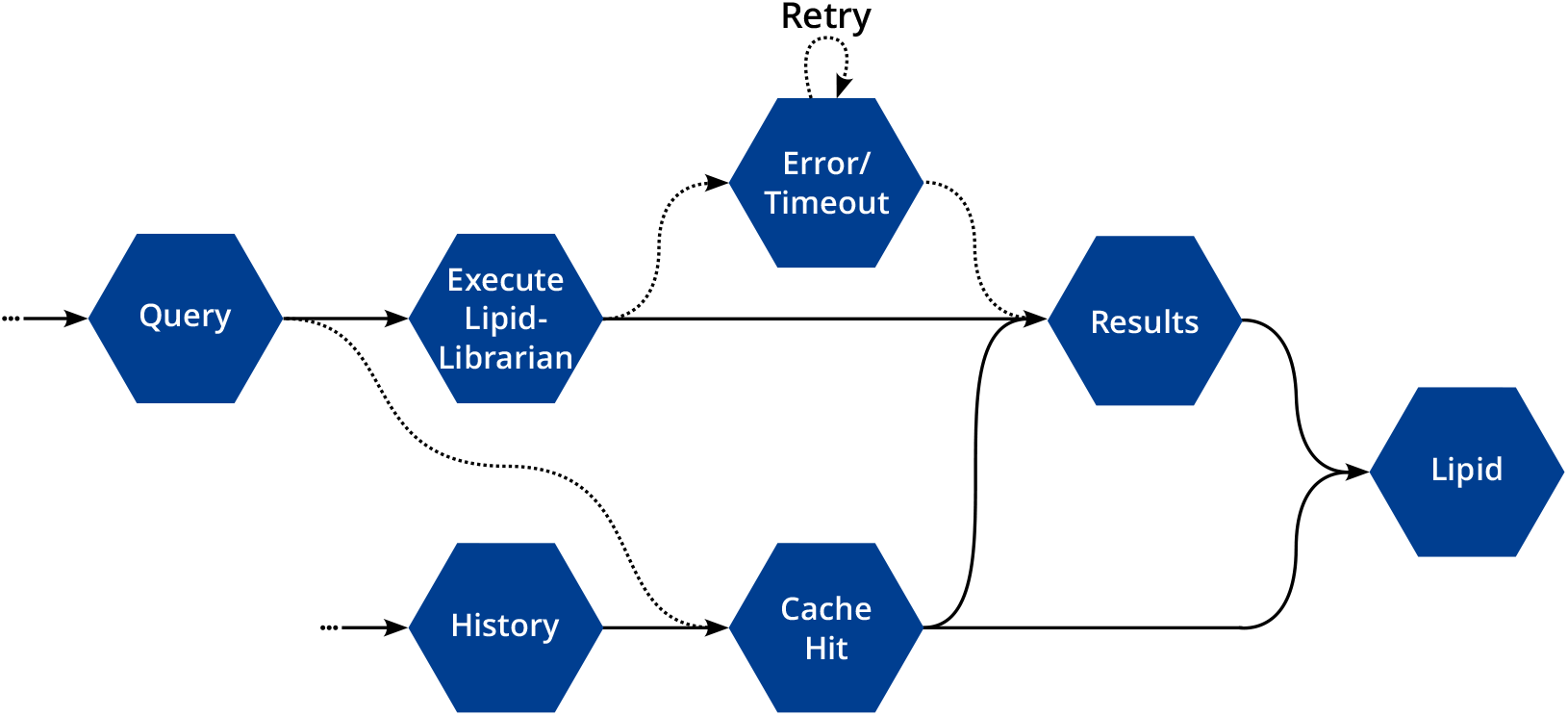
Conceptual workflow of LipidLibrarianWeb. The interaction with the frontend either starts with a query or with a revisit of a previous query via the history. If a query has already been executed (which is always the case if it accessed from the history), a cache hit skips the whole LipidLibrarian execution and leads directly to the results page or the lipid details page.

The query can lead to two possible results: Either an error occurs because the input is invalid, or a waiting screen appears while the user waits for the query to complete. While waiting for a result, a timeout may occur if a query to one of the databases takes too long to complete; in this case, the input is queried again. If the query executed successfully the results are saved into the database and will appear under the loading bar in the waiting screen. Alternatively the user can open the history side panel, pictured in Figure 4, by clicking on the history button in the navigation bar. Similar to the simple search field the history side panel can be accessed from every page of the website and enables the users to quickly inspect, edit and share the query results. At the bottom of the history panel (Figure 4, B) there is a session manager that enables users to access their old sessions or share their current session with co-workers or visit them on other devices. This is achieved by assigning a UUID session token to the query in a database, which can either be randomly generated or defined by the user, as long as it is in UUID format. Every visitor of the website automatically gets a token assigned if they accept the cookies from the website and execute a query, which they can find on the bottom of the history panel under current token. At the top of the panel (Figure 4, A), there is a list of all recently executed queries in chronological order with multiple lipids per query. If the user chooses to not accept the cookie with the session token, the history feature is unavailable, as there is no GDPR compliant way for us to know who they are if they refresh the page or close the browser window.

**Figure 4.**
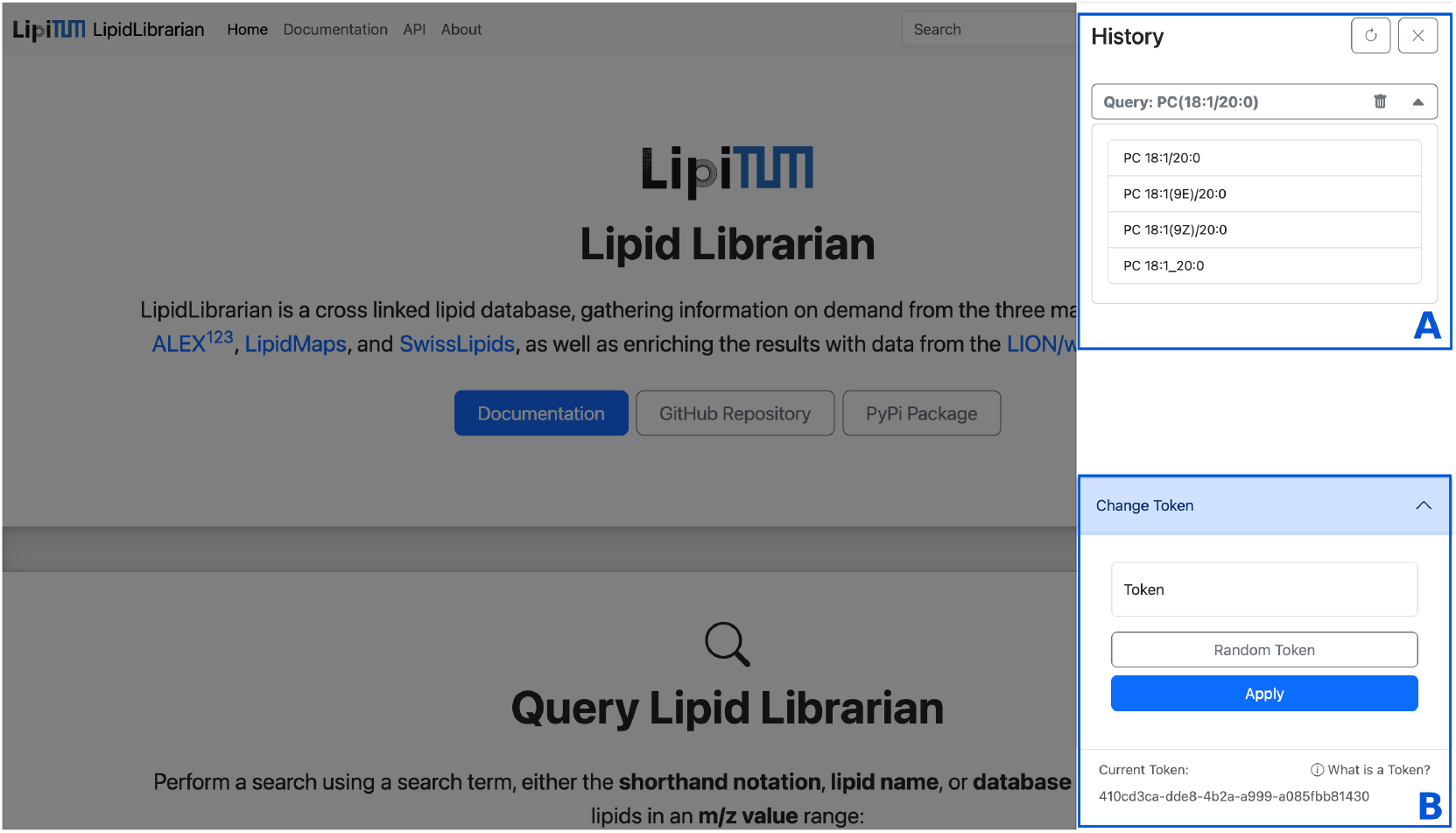
The home view of LipidLibrarianWeb with the history side panel open. In the upper part of the history (A) there is a list of all queries with their results, allowing the user to navigate quickly between previous queries. The history is bound to a token, which is displayed at the bottom of the panel (B). The user can share the token with another user or device to view and edit the associated history together.

In the final part of the workflow the user can select a lipid from the history panel and examine it in the results view, as shown in Figure 5. Here, the user can view all the information acquired about the selected lipid, starting with its name, hierarchy level, class, masses and information on the database sources from which LipidLibrarian has obtained information for this particular lipid. In addition, the results view provides details into adducts, fragments, database identifiers, structure identifiers, synonyms, ontology information and reactions associated with the lipid. The user can easily navigate and find information on each of these categories by using a navigation bar on the left side of the results.

**Figure 5.**
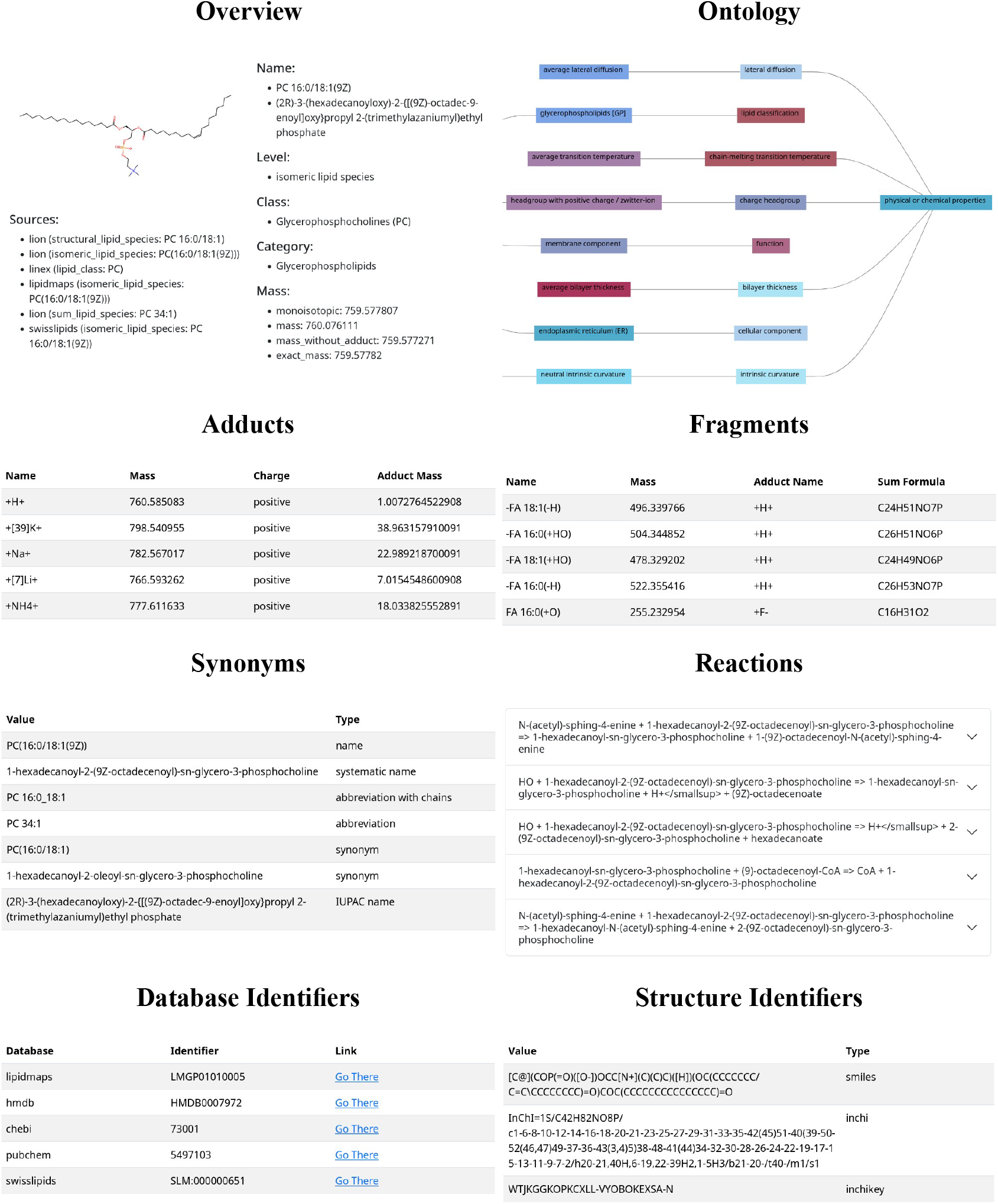
The results page of LipidLibrarianWeb divided into the individual categories: an overview over the lipid name, level, class, category and mass, its ontology, adducts, synonyms, fragments, reactions, database identifiers, and structure identifiers.

## 4 Discussion

LipidLibrarian implements the FAIR principles of findability, accessibility, interoperability, and reusability [31]. It is open source and depends only on other open source projects, shares data in a text-based JSON for machines to interpret and HTML for humans to read. It makes existing data, ideally from FAIR data sources, more findable, and keeps the data’s source always connected to the information for reference and attribution. The REST API is documented with the OpenAPI 3.0 standard. We also created LipidLibrarian with the idea in mind to integrate and therefore archive abandoned and defunct data sources and access the information connected to specific lipids and lipid classes. For example, ALEX^123^’s web server is not suitable for dynamic access via an API, which is why we extracted the data and created a way to make the content systematically accessible and searchable without being dependent on the web server.

From a technical point of view, we have ensured that the codebase is easy to maintain and service. For the end user, we have also ensured that the requirements for using LipidLibrarian are reasonable. All dependencies of the applications are available in PyPi and NPM, the standard repositories for Python and JavaScript, except for LipidLynxX, which we download, modify and build from its source code repository. For users who are still unable to compile or run LipidLibrarian, we provide alternative distribution options that are largely platform independent. We provide LipidLibrarian as a command line application and web service in OCI compatible containers and networks so that it is easy to install and use with software such as Docker, Podman or Kubernetes on almost any platform that supports them. The most critical part of LipidLibrarian, the parsing and conversion of lipid names, has been implemented redundantly with either LipidLynxX or Goslin, so if one of the two tools fails or is no longer available, our library can switch transparently between the two.

The LipidLibrarian library is designed to be easily extensible for new databases and resources. For databases and FAIR data sources and tools this is no issue, however, there are a few things to consider when including tools in LipidLibrarian. These include runtime, resource intensity and implementation complexity, which affects stability and maintainability. Especially API requests over the network that calculate complex results may take a lot of time in comparison to locally executing sub-processes that extrapolate additional information from efficiently stored data. Generally the optimal way to include a tool would be to import the external code written in the same language as LipidLibrarian to minimize the complexity needed for the code wrapping the execution, error handling and result interpretation, as well as the need for binaries or installation of other programming languages in the execution environment.

In the future, we plan to improve the *m/z* ratio query process, as the online databases do not provide sufficiently fast and relevant results from our experience. We also want to introduce a new publicly availabe API endpoint which allows to automatically start querying and redirecting to the most relevant lipid in a single request, so other tools and resources can link to LipidLibrarianWeb.

## 5 Conclusion

Lipidomics is a vast field with many undiscovered mysteries that we hope to make explorable and accessible with the introduction of LipidLibrarian. As the field grows, we expect more resources to be available. These resources may contain information that we have already presented here, like synonyms, classification, or mass. However, they could also offer novel information about lipids, such as their interactions with other common molecules. Regardless of the source or data type, LipidLibrarian is prepared to incorporate them.

In summary, LipidLibrarian enables effortless, comprehensive, and automated searching for lipid information, and enables a harmonized way to query information from the most important lipid databases as well as from archived abandoned data sources. By preventing tedious research, comparison, and aggregation of lipid information, it accelerates the research workflow, making it a meaningful tool for the scientific community.

## Data Availability Statement

All data used in the experiment is publicly available from the cited articles.

The LipidLibrarianWeb application is currently accessible under the URL https://lipidlibrarianweb.tools.lipidomics.dev.

## Code Availability Statement

Source code of the LipidLibrarian: GitHub (GNU AGPL v3 License): https://github.com/lipitum/lipidlibrarian

Source code of the LipidLibrarianWeb Application: GitHub (GNU AGPL v3 License): https://github.com/lipitum/lipidlibrarianweb

The LipidLibrarian Python package on PyPI: https://pypi.org/project/lipidlibrarian

## Author Contributions

JKP conceived the original idea and led the initial design of the project along with VW and NK. FN, KM, and KP developed the first version of the Python package with regular feedback from VW and NK. KP performed factual accuracy validation, conducted performance benchmarking, and carried out runtime analysis to ensure robustness and efficiency for the first version of the Python package. FN refactored and expanded the Python package to enhance its performance and usability. FN and KM wrote the script and database to extract and store MS/MS fragment data from ALEX^123^. FN and KM implemented the REST API to facilitate programmatic access to the tool. FN designed and developed the web frontend, KM contributed to the frontend implementation. JKP supervised the project and provided critical feedback and guidance throughout all stages. All authors contributed to writing and reviewing the manuscript and approved the final version.

## Acknowledgments

This project was funded by the Bavarian State Ministry of Science and the Arts in the framework of the Bavarian Research Institute for Digital Transformation (bidt; LipiTUM). https://www.bidt.digital/person/lipitum

